# Acoustic-based model estimation of snapping shrimp populations and the effects of a sponge die-off

**DOI:** 10.1101/056986

**Authors:** Jack Butler, Mark J Butler, Holly Gaff

## Abstract

Human use of the ocean and its ecosystems continues to degrade coastal habitats around the world. Assessing anthropogenic impacts on these environments can be cost and manpower intensive; thus, developing rapid, remote techniques to assess habitat quality has become increasingly important. We employed autonomous hydrophone receivers to record the soundscapes of healthy hard-bottom habitat in Florida Bay, Florida (USA) and hard-bottom areas impacted by sponge die-offs. We also recorded sounds emanating from individual sponges of three species that were isolated in underwater sound booths, and then enumerated the invertebrates (mostly snapping shrimp) dwelling within the canals of each sponge. From these recordings, a modified cylindrical sound propagation model was used to estimate distances to snapping shrimp snaps. Using the program *Distance*, which applies distance sampling theory to cue count surveys, we estimated snapping shrimp population density and abundance within both habitat types. More snapping shrimp snaps per unit time were recorded in healthy hard-bottom areas as compared to degraded hard-bottom areas. In addition, the average distance to a snap source was greater within degraded hard-bottom areas than within healthy hard-bottom areas. As a consequence, the estimated density and abundance of snapping shrimp were one to two orders of magnitude greater within healthy habitat than within degraded habitat. This study demonstrates the feasibility of using acoustic sampling and modeling to rapidly assess populations of soniferous benthic indicator species, whose vocalizations may yield indirect estimates of habitat quality.

## 1. Introduction

Humans rely upon ocean ecosystems for goods and services; unfortunately, extractive use of marine resources (e.g., fishing and mining,) and the indirect effects of human habitation (e.g., land-based run-off and climate change) have altered and degraded these ecosystems (Jackson et al. 2001; Halpern et al. 2008). Worldwide, marine ecosystems are declining (Suchanek 1994; Valiela et al. 2001; Waycott et al. 2009) and coastal ecosystems are particularly vulnerable to anthropogenic disturbances (Vitousek et al. 1997; Limburg 1999; Lotze & Milewski 2004), threatening their function (Solan et al. 2004; Worm et al. 2006; Diaz & Rosenberg 2008).

Habitat monitoring and assessment are key to understanding how ecological communities respond to habitat degradation (Kremen et al. 1994), yet monitoring presents many challenges. It is often time-consuming, expensive (Harris et al. 2015), and prone to human bias (Willis 2001); as when, for example, the avoidance of divers by fishes skews estimates of their biodiversity (Dickens et al. 2011). So the development of accurate and inexpensive monitoring techniques is becoming increasingly important as anthropogenic influences continue to buffet near-shore environments, including structurally complex coastal habitats that are so important as nurseries and foraging grounds (Airoldi et al. 2008).

One promising technique, based on the burgeoning science of soundscape ecology, relies on the measurement of sound to monitor ecosystems (Pijanowski et al. 2011). Although pioneered in terrestrial ecosystems, the study of soundscapes has been extended to the marine environment as a framework for environmental monitoring (Harris et al. 2015). Contrary to the public perception that the sea is a quiet realm-as implied, for example, in Jacques Cousteau’s *Silent World*-the ocean is alive with sound.

Underwater sound, whose sources are physical, biological, and anthropogenic, has been studied for decades. Early studies by Tait (1962) and Cato (1976, 1980) were some of the first to describe variation in underwater noise from rock and coral reefs off New Zealand and Australia. Recent research has confirmed that many of those noises are of biological origin and exhibit diel, lunar, and seasonal variation (Radford et al. 2008a, 2008b). There is also spatial variability in the sounds that emanate from within and among habitats (Radford et al. 2008a, 2008b, 2010; Lillis et al. 2014), but only a few studies have used acoustics to assess community structure or habitat characteristics in the marine environment. For example, Lammers et al. (2007) described how acoustic activity is correlated with the structural characteristics of habitats, whereas Kennedy et al. (2010) determined that acoustic variability was positively correlated with the density, biomass, and diversity of organisms on coral reefs.

Though many marine organisms produce sounds and contribute to the biological component of soundscapes (Myrberg 1981; Versluis et al. 2000;Bouwma & Herrnkind 2009; Schärer et al. 2014; Staaterman et al. 2014), few are as ubiquitous as snapping shrimps whose snaps contribute a significant portion of energy to the biological din (Au & Banks 1998; Radford et al. 2008a, 2010; Bohnenstiehl et al. 2016). By rapidly closing the dactyl of its enlarged chela, a snapping shrimp creates a cavitation bubble that produces a loud pop upon its collapse (Versluis et al. 2000). Snapping shrimps occur throughout temperate and tropical waters (Au & Banks 1998; Cato & McCauley 2002; Radford et al. 2010) and dwell in a variety of habitats, from estuaries to coral reefs (Au & Banks 1998). One group of snapping shrimps within the genus *Synalpheus*, a clade of ~ 100 species, all live within the canals of tropical sponges (Duffy & Macdonald 1999; Duffy 2002). Some species of *Synalpheus* live in colonies of several hundred shrimps all living within the same sponge; a few have developed eusociality, the only occurrence of this extreme form of social behavior known among marine animals. Many species within this genus exhibit direct development in which eggs hatch directly into crawling juveniles (Duffy 2002), which further reinforces the link between shrimps and their sponge home.

Large sponges that harbor snapping shrimps are particularly abundant and important components of tropical hard-bottom communities, such as those found in the Florida Keys (USA). Hard-bottom habitat covers roughly 30% of the near-shore environment of the Florida Keys where dozens of sponge species dominate the benthic animal biomass with a mean density of >80,000/ha (Stevely et al. 2011). Many of those sponges, especially large sponges like the loggerhead sponge (*Spheciospongia vesparium*), provide shelter and habitat for fish and invertebrates (Butler et al. 1995), including those that are soniferous (i.e., “sound producers”).

However, the Florida Keys have undergone drastic ecological change in recent decades. In 1991, 2007, and 2013 portions of the Florida Keys-especially Florida Bay, the bay lying between the Florida mainland and the islands of the Florida Keys-were subjected to prolonged thermal stress and a major shift in salinity due to anomalous and persistent weather conditions (Butler et al. 1995; Stevely et al. 2011). These physical stresses resulted in massive and widespread blooms of cyanobacteria whose radical increase in concentration precipitated the mass mortality of sponges within a 500-km^2^ area of the bay (Butler et al. 1995). The widespread loss of sponges resulted in a significant reduction of structural complexity in affected areas, leaving barren expanses of open substrate where sponges were once numerous. The ecological effects of such a dramatic shift in the character of these systems are still being studied, among these being a significant change in the underwater acoustic signature of affected hard-bottom areas (Butler et al. 2016).

Because of the close association between snapping shrimps and sponges, a reduction in sponge density in places such as Florida Bay would also likely reduce snapping shrimp density and abundance. Thus, the present study aimed to: (1) evaluate the efficacy of using remote acoustic monitoring to estimate snapping shrimp density and abundance, and (2) examine how sponge mortality might have affected the distribution of snapping shrimp populations in Florida Bay.

## 2. Materials and methods

To evaluate the effect of loss of sponges on snapping shrimp populations, underwater soundscapes were recorded at six healthy hard-bottom sites in Florida Bay outside of the area impacted by the sponge die-offs and at five hard-bottom sites within the area affected by the sponge die-off (Fig. 1). Butler et al. (2016) determined that the number of snapping shrimp snaps produced in hard-bottom areas unaffected by sponge die-offs was greater than the number of snapping shrimp snaps produced in hard-bottom areas degraded by sponge die-offs. Therefore, acoustic recordings made outside the range of the sponge die-offs were used as a baseline to characterize the soundscapes indicative of healthy snapping shrimp populations to which recordings made within the die-off area were compared.

### 2.1. Acoustic analysis and number of snapping shrimp snaps

Habitat recordings were made using submersible hydrophone (Fig. 2A). Each system consisted of a manufacturer-calibrated Aquarian Audio H2a omnidirectional hydrophone (Aquarian Audio Products: sensitivity-180 dB re 1V/uPa) connected to a Roland R-05 solid-state WAV recorder (48 kHz, 16 bit) housed within a waterproof housing. The system was manually calibrated using pure sine signals from a signal generator, measured in line with an oscilloscope, and the recordings were analyzed in MATLAB 2014b software (Mathworks, Inc.) by code written specifically for the calibration of hydrophone systems. Fifteen-minute recordings were made at noon during either the first quarter or last quarter moon phase at each site, and from these recordings five 10-second subsamples were extracted for further analysis. All recordings were post-processed using MATLAB 2014b software (Mathworks, Inc.). Each 10-second subsample was processed through a MATLAB script written specifically for this study. First, the data were high-pass filtered to 100 Hz to remove extraneous low-frequency interference. The data were then plotted for visual inspection and to determine a snap count threshold level. The threshold is the level above which any transient spike in the data is considered a snapping shrimp “snap” and varies from recording to recording. Using the threshold level, data for individual snaps within the recording were located, extracted, and stored. Once data for individual snaps within a given subsample were extracted, the peak-to-peak pressure level for each snap was calculated. A calibration factor was applied to the raw data to calculate absolute
135 pressure levels, which were converted to decibels relative to 1 microPascal (dB re 1 μPa). These values were later used to calculate sound transmission loss, as described below.

**Figure 1.**
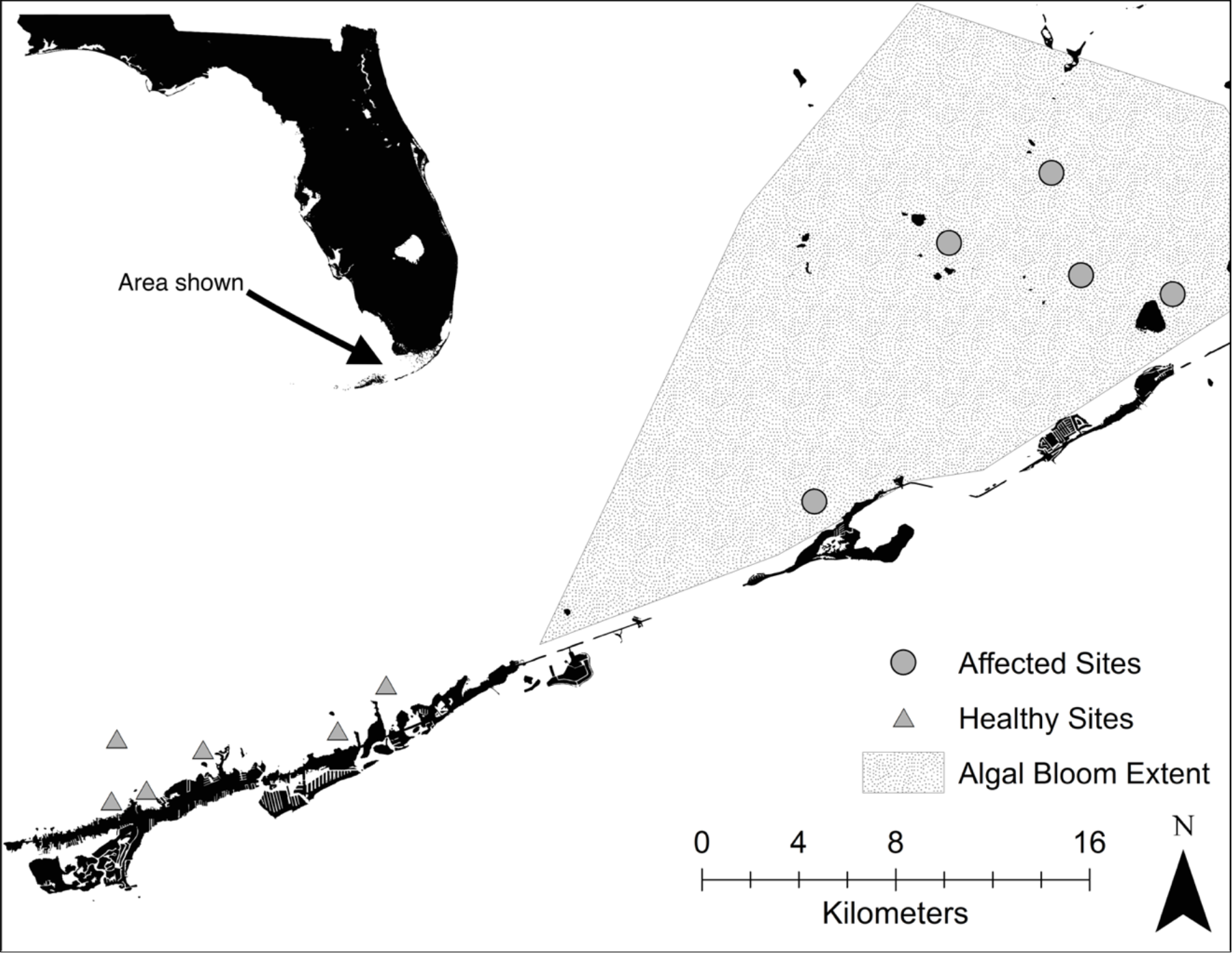
Map of study area showing algal bloom extent and acoustic recording sites

**Figure 2.**
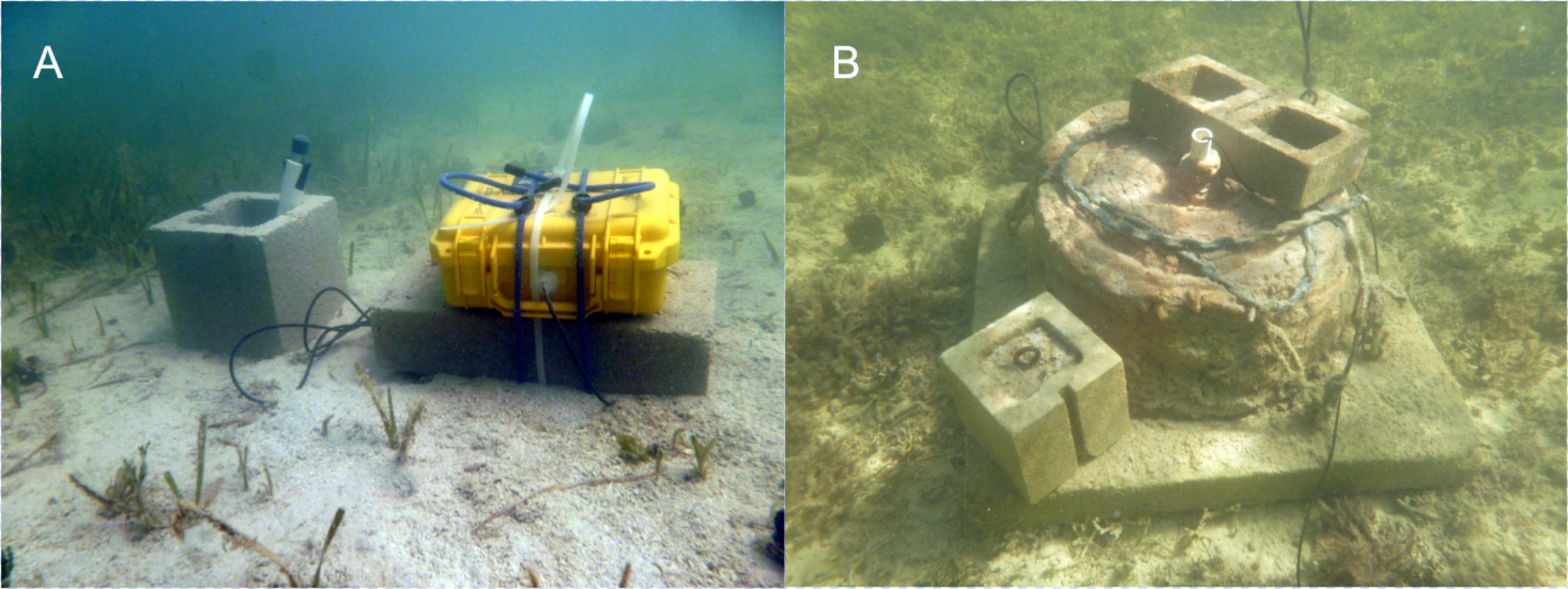
A remote hydrophone system (A), and the in situ sound damping chamber (B)

### 2.2. Estimating snapping shrimp snap rate and snap source level

To determine the cue rate (i.e., snap rate) and to estimate the snap source level, 15 individual sponges of three sponge species (loggerhead sponge, *Spheciospongia vesparium*; sheepswool sponge, *Hypospongia lachne*; yellow sponge, *Spongia barbara*) in which snapping shrimps can be found were acoustically isolated *in situ* using an underwater sound damping chamber. The chamber was constructed of a tin washtub (54 cm dia; 26 cm ht) encapsulated with 5 cm of closed-cell foam and set in a 15 cm thick concrete base to render it negatively buoyant (Fig. 2B). A hydrophone attached to a WAV recorder (see description above) was lowered through a 3 cm dia tube at the center of the chamber, permitting the recording of sound from individual sponges *in situ*.

The effectiveness of the sound-damping chamber was tested in two ways. First, simultaneous *in situ* recordings of the soundscape acoustic spectra outside the chamber were compared to the acoustic spectra within the chamber (Fig. 3A). In addition, after recording the number of snapping shrimp snaps as described above, only snaps whose power exceeded a threshold that excluded quieter snaps recorded outside the sound-damping chamber were counted (Fig. 3B). This was done to ensure the chamber was effectively quieting snaps external to the chamber to reduce false snap counts from sponges recorded within the chamber. Once the chamber was positioned over the sponge, sound levels were recorded for 15 minutes. The sponge was then cut from the substrate and placed in a plastic bag for transport to the laboratory where it was dissected so as to remove and count all of the infaunal organisms, which were primarily snapping shrimps.

For recordings of sponges that housed snapping shrimps, the number of snapping shrimp snaps emanating from that single sponge was calculated as described above. Using the number of snaps and the number of snapping shrimp found within each sponge, cue rate (snaps/10-sec/shrimp) was calculated. For each acoustically-isolated sponge, a calibration factor was applied to the recording and the peak-to-peak sound pressure level (dB re 1 μPa) for individual snaps was calculated. The snap source levels for all individual snaps were averaged to determine the source level used to calculate transmission loss.

### 2.3. Estimating snapping shrimp distance to hydrophone receiver

For each 10-second subsample of a recording, the previously calculated sound source level (SL) and the received sound level (RL) for each snap (see above) was used to calculate the transmission loss of each snap. The transmission loss (TL) of each snap was calculated as:

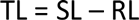

Using the transmission loss for each snap, the distance (d) of each sound source to the hydrophone receiver was calculated using a modified cylindrical spreading model. Sound propagates cylindrically in shallow water habitats like the hard-bottom habitats (< 2m depth) in which these recordings were made. A cylindrical spreading model predicts that the coefficient of transmission loss should be 10 (Urick 1983), but to account for sound wave interference, sound absorption and scattering at the sea floor, and sound scattering at the sea surface, the coefficient of transmission loss was raised to 15:

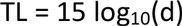

From this equation, the distance from the sound source to the hydrophone receiver was calculated:

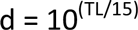

**Figure 3.**
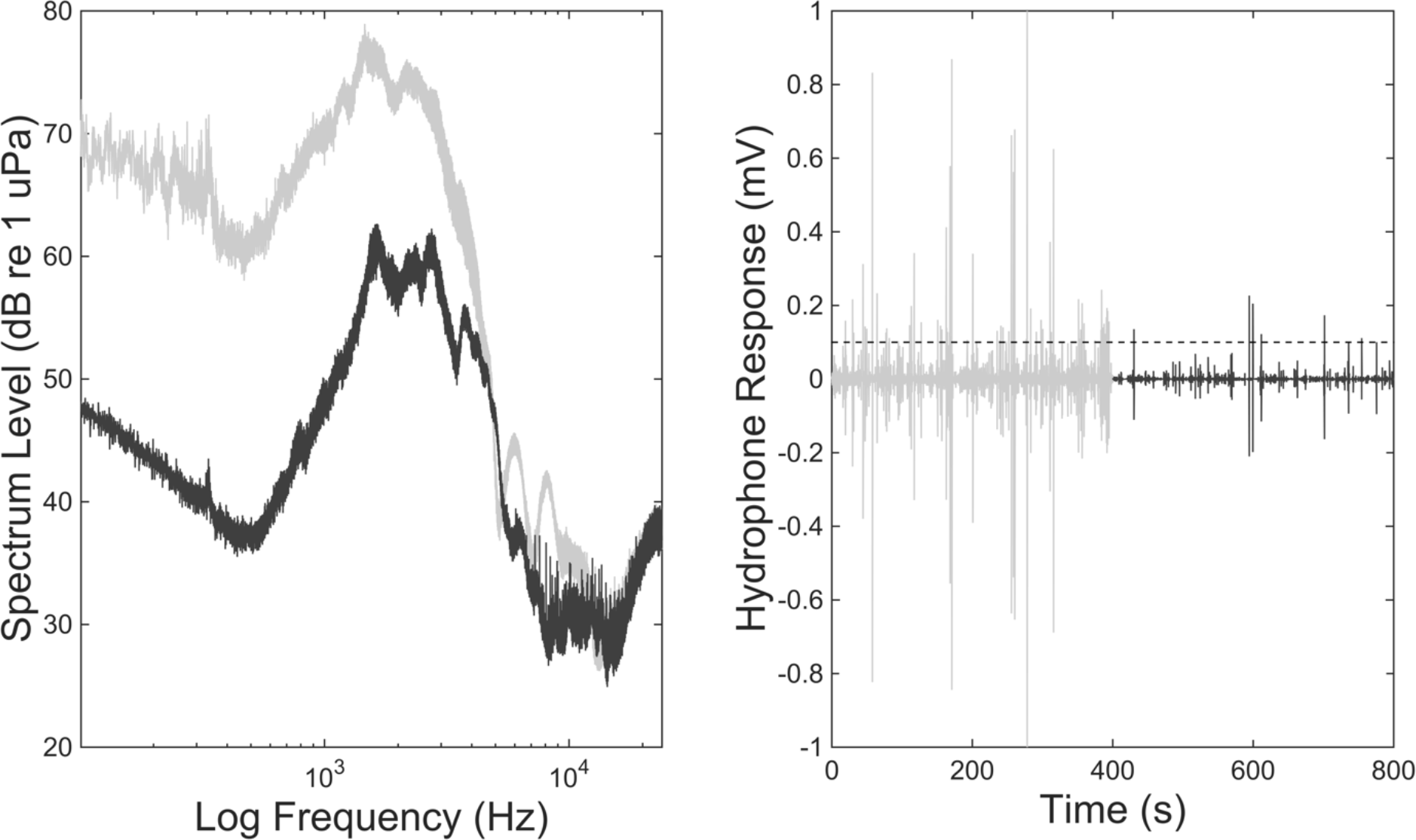
(A) shows the comparison between the acoustic spectra outside the sound damping chamber (gray) versus the acoustic spectra within the chamber (black). (B) shows the occurrence of snapping shrimp snaps above a set threshold level (dashed line) outside Snapping shrimp population density and abundance estimation.

### 2.4. Snapping shrimp population density and 195 abundance estimation

Using the snap rate calculated within the sound-damping chamber and the estimated distances to snapping shrimp snaps within each of the habitat recordings, snapping shrimp population density was estimated using the program *Distance* (version 6.2; Thomas et al. 2010). Distance sampling techniques for population density estimation are widely used (Buckland et al. 2005) and have recently been modified to suit point transects wherein distances to sound cues (e.g., bird calls or whale songs) are used rather than distances to animal sightings. This technique allows for easy and inexpensive remote sensing and density estimation of any soniferous organism and has been successfully implemented within the marine environment (e.g.,Küsel et al. 2011; Harris et al. 2013). Population density estimation via distance sampling uses a probability density function based on an underlying detection function; this detection function represents the probability of detecting a cue of interest given its distance from the receiver (Marques et al. 2013). It is assumed that all organisms (or cues) of interest that lie directly on the transect line or point are counted with certainty, and the probability of detection declines monotonically with increasing distance from the line or point transect. The distribution of observed detection distances is used to estimate the average probability of detection, and this in turn is used to estimate population density and abundance. In addition, the cue rate (i.e., how often one organism makes one cue of interest) is used to convert cue density and abundance to animal density and abundance (Marques et al. 2009) by multiplying the cue density estimation by the cue rate.

*Distance* fits several detection functions to the sampling data and uses AIC to determine which model best fits the data. Once a model is selected, *Distance* provides a summary of the analysis, including a density estimate and 95% confidence intervals around that estimation. Thomas et al. (2010) suggest a half-normal detection function with a cosine adjustment for most studies, and this detection function best fit the data from the present study. Distances were truncated to 50 meters (that is, any snap estimated to be over 50 meters from the hydrophone receiver was removed from the population density and abundance estimates-about 1% of distance estimates) to avoid adding extraneous adjustment terms to the underlying detection function (Buckland et al. 2001; Thomas et al. 2010). Because distances were truncated, density and abundance estimates of snapping shrimp populations produced by *Distance* are based on a circular area with a radius of 50 meters, thus resulting in a total coverage area of 7854 m^2^.

Snapping shrimp population density (shrimp/m^2^) was estimated for each site, and the difference in shrimp density between healthy hard-bottom sites and degraded hard-bottom sites was tested using a nested ANOVA (site nested within habitat type). Pooled data for all sites within either the healthy hard-bottom area or the degraded hard-bottom area were used to estimate population density for healthy and degraded areas. Upper and lower 95% confidence interval density estimates were multiplied by the coverage area to obtain upper and lower snapping shrimp abundance estimates. In addition, number of snapping shrimp snaps per 10-second subsample and average distance to snap source were analyzed using nested ANOVAs (site nested within habitat type) to determine differences in those estimates between healthy hard-bottom and degraded hard-bottom areas.

## RESULTS

### 3.1. Sponge infaunal shrimp communities, snap rate, and snap source level

At least one individual of each sponge species harbored snapping shrimp of the genus *Synalpheus*, but the occurrence and abundance of snapping shrimp varied among sponge species (Table 1). Every loggerhead sponge housed snapping shrimp in high abundance (mean number of shrimp per sponge: 28 ± 24.15 s.d.), and 80% of the sheepswool sponges housed snapping shrimps albeit at lower abundance (13.5 ± 14.65); however, only 20% of the yellow sponges housed any snapping shrimp, and then only in low numbers (1.69 ± 5.72). The number of snapping shrimp within a given loggerhead sponge scaled linearly with sponge volume (r^2^=0.703; p<0.001), but the relationship between sheepswool (r^2^=0.217; p = 0.244) and yellow sponge (r^2^=0.034; p = 0.91) volumes and number of snapping shrimp within each sponge were weak and non-significant (Fig. 4). However, these differences could be attributed to a wider size range for larger loggerhead sponges compared to sheepswool and yellow sponges, which are smaller with less size variability. The number of snaps per shrimp per ten-seconds was 0.014 ± 0.023 (mean ± s.d.), and the average peak-to-peak source level of all snaps was 130 dB re 1 μPa over 100-24,000 Hz.

**Figure 4.**
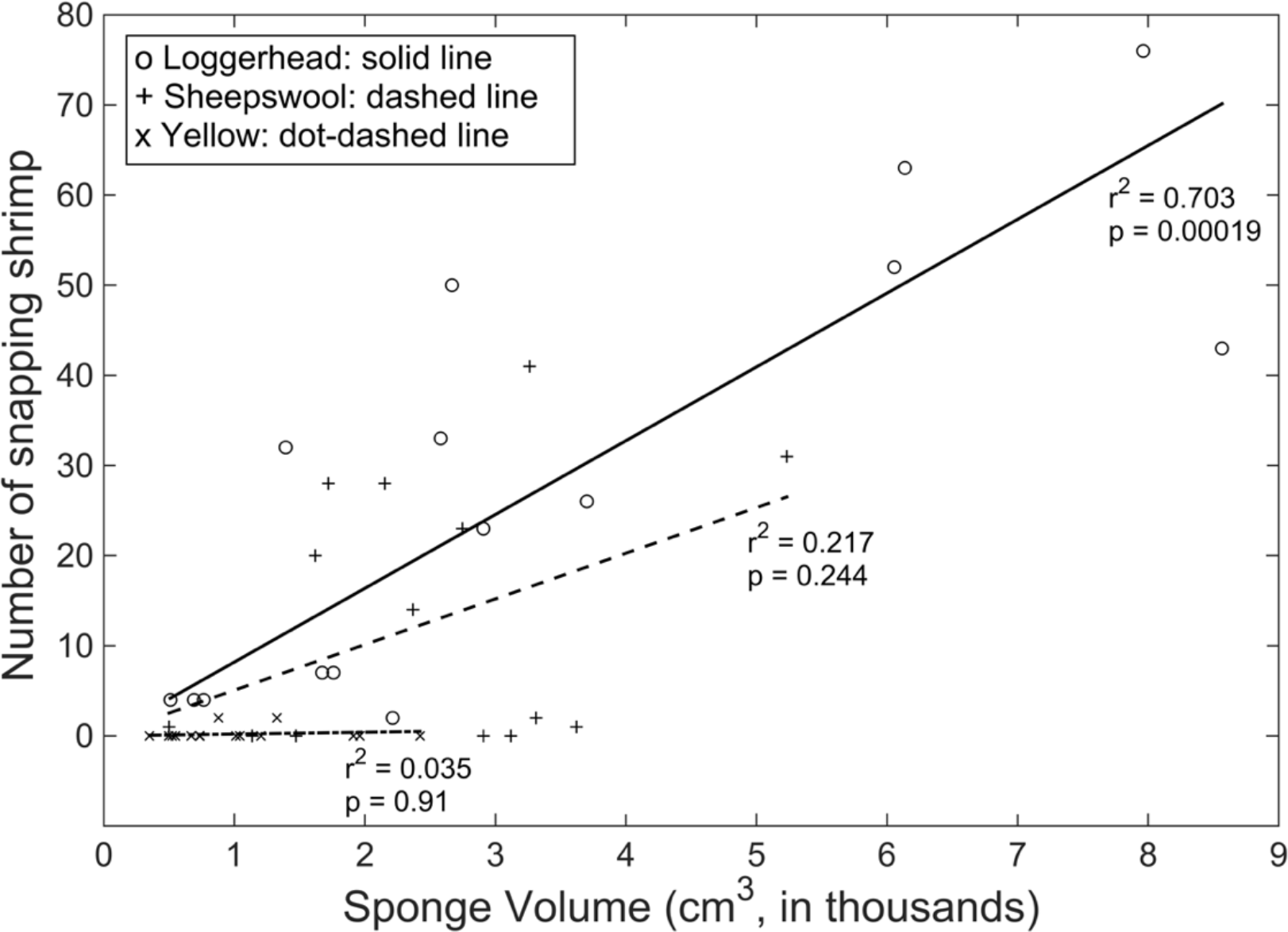
Sponge volume versus number of snapping shrimp found within individual sponges

**Table 1.**
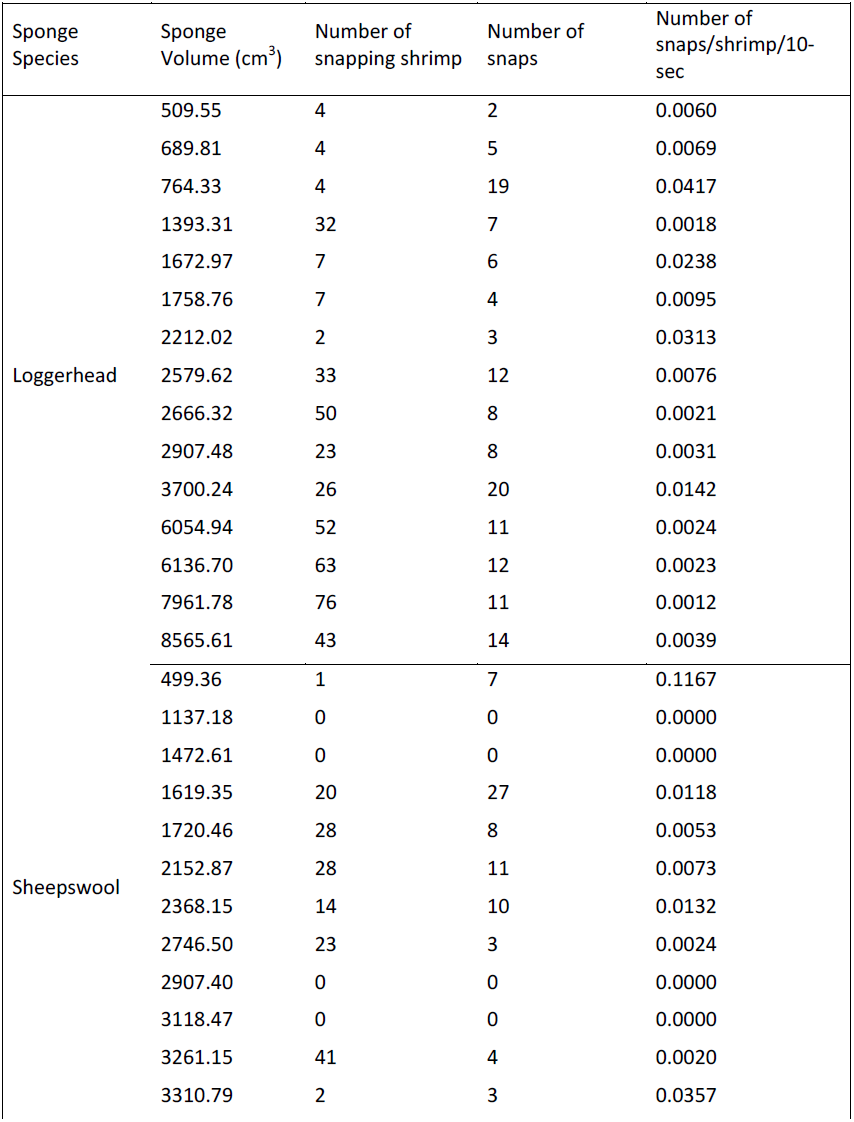
Summary of infaunal shrimp communities within individual sponges.

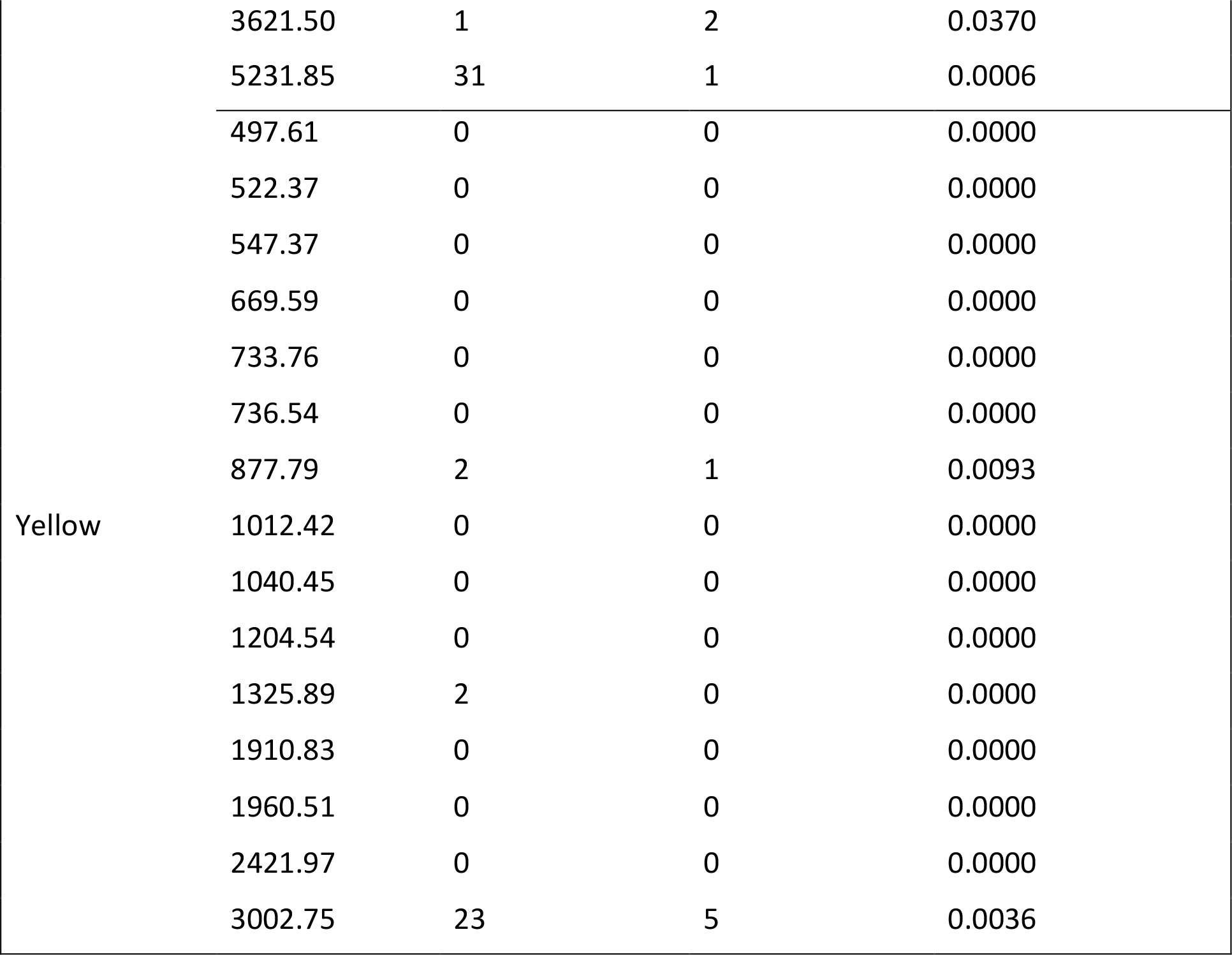

### 3.2. Number of snapping shrimp snaps and distance to snap source

Habitat type significantly affected the number of snapping shrimp snaps per ten-seconds, as well as the average distance to a snap’s source. The average number of snapping shrimp snaps per ten-seconds in healthy hard-bottom areas (241 ± 26.67; mean ± s.e.) was significantly greater (F_1,9_ = 72.9, p < 0.001) than the average number of snapping shrimp snaps per ten-seconds in affected hard-bottom areas (35 ± 5.98). Conversely, the average distance to a snap’s source in degraded hard-bottom areas (18.24 m ± 0.86; mean ± s.e.) was significantlygreater (F,_1,9_ = 57.38, p ≪ 0.001) than the average distance to a snap’s source in healthy hard-bottom habitat (6.94 m ± 0.15).

### 3.3. Snapping shrimp population density and abundance estimation

Snapping shrimp population density estimates also differed significantly among habitat types (F_1,9_ = 13.84, p = 0.002); those within healthy hard-bottom areas (2.68 shrimp per m^2^ ± 0.68; mean ± s.e.) were greater than estimates of shrimp density within degraded hard-bottom areas (0.057 ± 0.013). Abundance estimates varied among sites, but degraded sites exhibited lower abundance estimates than did healthy sites (Table 2). The lowest abundance estimated at degraded sites was 23 shrimp within the hydrophone coverage area (7854 m^2^), and the highest snapping shrimp abundance estimated at degraded sites was 7,657 shrimp within the hydrophone coverage area. The estimated abundance of snapping shrimp within healthy sites was one to two orders of magnitude greater than estimates within degraded sites. The lowest estimate within healthy sites was 471 shrimp, and the highest estimate was 341,248 shrimp within the hydrophone coverage area.

**Table 2.**
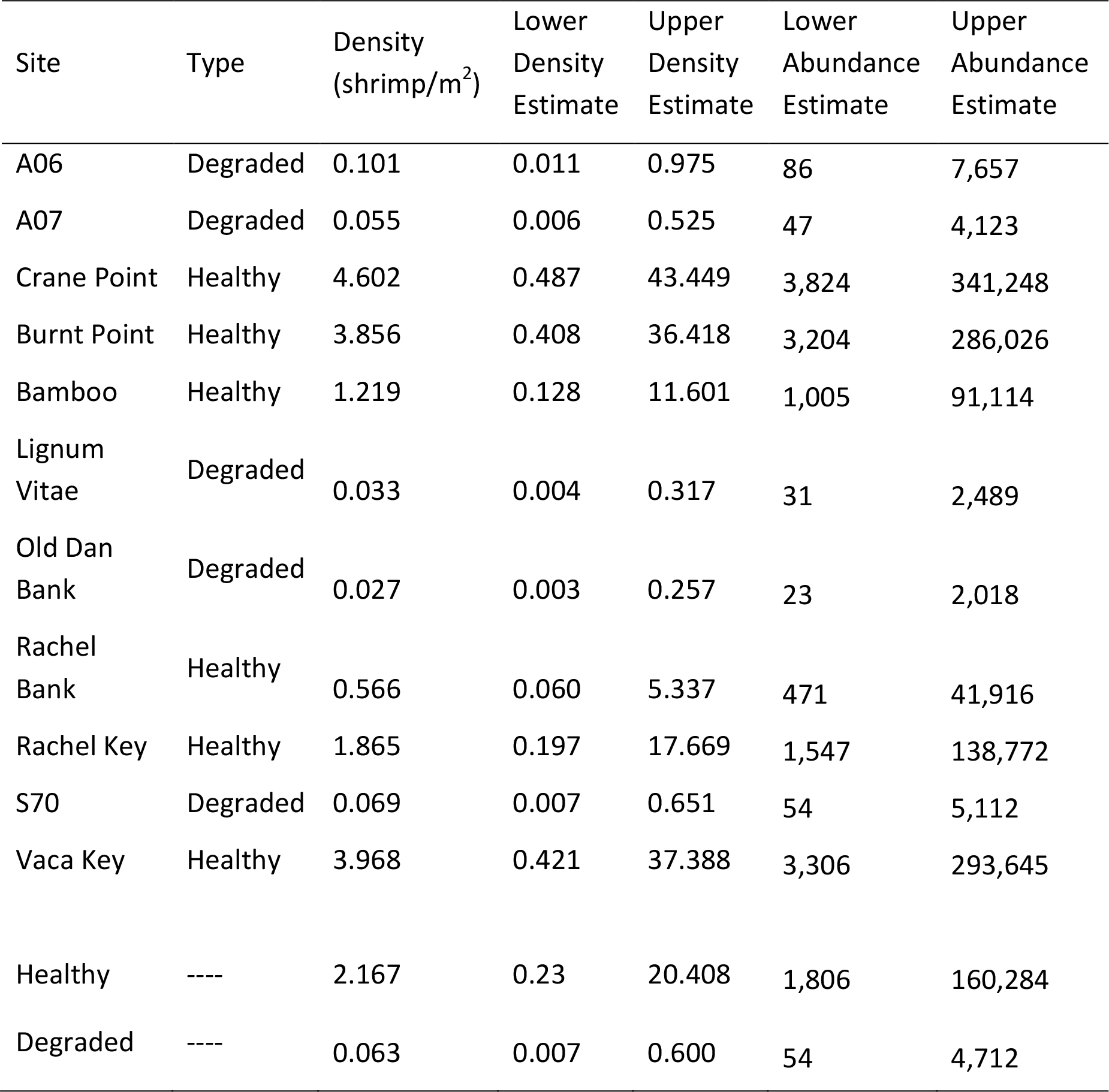
Estimated snapping shrimp population densities and abundances.

| Site | Type | Density<br>(shrimp/m<sup>2</sup>) | Lower<br>Density<br>Estimate | Upper<br>Density<br>Estimate | Lower<br>Abundance<br>Estimate | Upper<br>Abundance<br>Estimate |
| --- | --- | --- | --- | --- | --- | --- |
| A06 | Degraded | 0.101 | 0.011 | 0.975 | 86 | 7,657 |
| A07 | Degraded | 0.055 | 0.006 | 0.525 | 47 | 4,123 |
| Crane Point | Healthy | 4.602 | 0.487 | 43.449 | 3,824 | 341,248 |
| Burnt Point | Healthy | 3.856 | 0.408 | 36.418 | 3,204 | 286,026 |
| Bamboo | Healthy | 1.219 | 0.128 | 11.601 | 1,005 | 91,114 |
| Lignum<br>Vitae | Degraded | 0.033 | 0.004 | 0.317 | 31 | 2,489 |
| Old Dan<br>Bank | Degraded | 0.027 | 0.003 | 0.257 | 23 | 2,018 |
| Rachel<br>Bank | Healthy | 0.566 | 0.060 | 5.337 | 471 | 41,916 |
| Rachel Key | Healthy | 1.865 | 0.197 | 17.669 | 1,547 | 138,772 |
| S70 | Degraded | 0.069 | 0.007 | 0.651 | 54 | 5,112 |
| Vaca Key | Healthy | 3.968 | 0.421 | 37.388 | 3,306 | 293,645 |
| Healthy | ---- | 2.167 | 0.23 | 20.408 | 1,806 | 160,284 |
| Degraded | ---- | 0.063 | 0.007 | 0.600 | 54 | 4,712 |

## 4. Discussion

Habitat degradation disproportionately affects coastal marine ecosystems (Vitousek et al. 1997; Limburg 1999; Lotze & Milewski 2004), and Florida Bay, where this study took place, is no different. Extensive, persistent cyanobacteria blooms have repeatedly decimated hard-bottom sponge communities (Butler et al. 1995; Stevely et al. 2011), resulting in markedly different soundscapes including fewer snapping shrimp snaps within degraded hard-bottom areas (Butler et al. 2016). The present study confirms that there are indeed fewer snapping shrimp snaps per ten-second subsample at degraded sites. We also estimated distances among individual snaps and found that the distance to a snap’s source (i.e., the distance from a snapping shrimp to the hydrophone) was greater on degraded sites than on healthy sites. This increase in distance to a snap source coincides with a reduction in the number of large, canal-bearing sponges in which the majority of snapping shrimp can be found. Furthermore, shrimp abundances estimated using distance-sampling techniques indicate that degraded hard-bottom sites harbor fewer snapping shrimp, and that these populations are less dense.

Many snapping shrimp species within the genus *Synalpheus* dwell commensally within a diverse array of tropical and sub-tropical sponge species (Duffy 1992). However, living within sponges has some consequences on the shrimps, most notable is a limitation on shrimp size imposed by the canal structure of the host sponges. Duffy (1992) found no evidence of shrimp-induced canal excavation among the four tropical sponge species in which snapping shrimp were found (including the loggerhead sponge), and thus concluded that shrimp size must be limited by the canal size of the host sponge. Of the four sponge species Duffy (1992) dissected, the loggerhead sponge exhibited the greatest variability of canal widths (Figure 1 in Duffy 1992). Though we did not measure the canal structure of the three sponge species we dissected for this study, we observed that the loggerhead sponges had more well-developed canals compared to either the Sheepswool sponges or the Yellow sponges. This difference in canal structure could likely affect the numbers of snapping shrimp found within individual sponges (Table 1 & Fig. 4).

### 4.1. Passive acoustics and distance sampling as monitoring tools

As the world’s ocean ecosystems-particularly coastal ecosystems-continue to degrade (Jackson et al. 2001; Lotze & Milewski 2004), effective monitoring to determine the health of a target population or habitat is becoming increasingly important (Kremen et al 1994; Watanabe et al. 2002; Airoldi et al. 2008). Passive acoustic monitoring provides a suite of tools to answer scientific and management questions and avoids many downfalls of other surveying methods (Willis 2001; Dickens et al. 2011; Harris et al. 2015).

Monitoring of marine mammals and fishes via passive acoustics has increased in recent years (Moore et al. 2006; Mellinger et al. 2007; van Opzeeland et al. 2008; Luczkovich et al. 2008). Many species of marine mammals and fishes produce distinct sounds. For example, whales are often not amenable to visual survey methods, but can be monitored using passive acoustics. Antarctic blue whales produce characteristic long (~20 sec) tonal calls that downsweep from 28Hz to 18Hz (Sirović et al. 2004, 2009), whereas Antarctic fin whales produce short (~1 sec) calls downswept from 28Hz to 15Hz, often with a short following call around 89Hz (Sirović et al. 2004). Combined visual and passive acoustic surveys have shown that the passive acoustic surveying techniques detect as many as ten times more cetaceans than the visual surveys (McDonald & Moore 2002; Sirović et al. 2004; Barlow & Taylor 2005). In addition, passive acoustic monitoring can continue throughout the night and through conditions that would make visual surveys impossible (Mellinger & Barlow 2003; Mellinger et al. 2007).

However, if passive acoustic monitoring is to serve as a robust monitoring technique, it must also provide information such as population density or abundance. Several studies on cetacean abundance have combined line-transect surveys and passive acoustics to estimate the population density of the species-of-interest within a given geographic area (McDonald & Fox 1999; Barlow & Taylor 2005; Moretti et al. 2006; Mellinger et al. 2004; Marques et al. 2009). Marques et al. (2009) extended distance sampling techniques (i.e., estimating a detection function based on an animal-of-interest’s distance to a point or line transect) to point transects of cue counts of Blainville’s beaked whale (*Mesoplodon densirostris*) and estimated the detection function using digital acoustic tag (DTag) data from a previous study (Johnson & Tyack 2003). Extending the work of Marques et al. (2009), Küsel et al. (2011) estimated the population density of Blainville’s beaked whale using a single fixed hydrophone rather than an array of hydrophones and estimated distance to each whale call from the hydrophone by estimating the detection function and enlisting the passive sonar equation (equation 2 in Küsel et al. 2011). The density estimate of beaked whales in the Küsel et al. (2011) study was nearly three times higher than the estimate of Marques et al. (2009), however, re-analysis of the Marques et al. (2009) estimate based on one hydrophone (rather than an array of 82, as was in the original study) brought the two estimates closer together.

Similar to Küsel et al. (2011), we used the passive sonar equation to estimate distances to the source of each snapping shrimp snap. Unlike Küsel et al. (2011), who had available previous estimates of beaked whale population density based on an accurate method of estimation (Marques et al. 2009) with which they could compare their methods, we had no comparable data on snapping shrimp density with which to compare our estimates. To estimate distances to snap sources more accurately, and thus estimate the detection function more accurately, we could in the future employ hyperbolic localization techniques. For that method, multiple hydrophones are deployed in an array and arrival-time differences of a sound of interest are determined among hydrophones so as to more precisely ascertain the sound source (Spiesberger & Fristrup 1990; Spiesberger 1999, 2001). In this study we used a single hydrophone receiver at each site and the passive sonar equation to estimate distance to a sound source. We could not localize that source, which would have been possible with an array of four or more hydrophones and hyperbolic localization techniques. However, the cost of obtaining and deploying a multiple hydrophone array was beyond the limits of the present study.

Because of the association between snapping shrimp and sponges on hard-bottom communities, employing multi-hydrophone arrays and hyperbolic localization techniques to find clusters of snap sources and mapping these resultant clusters onto a 2-D coordinate system-rather than just counting snaps-would allow for a conservative estimation of sponge biomass and location within a hard-bottom area. For example, using the shrimp number and sponge volume data collected while determining cue rate and fitting a simple linear model, we can predict sponge volume (a proxy for sponge biomass) on a given site. This method might work particularly well for estimating loggerhead sponge biomass, given the significant positive relationship between loggerhead sponge volume and number of snapping shrimp found within a sponge (Fig. 4), as well as the significant positive relationship between total loggerhead sponge volume on a given site and the number of snapping shrimp snaps produced on that site (Fig. 5). These predictions would, obviously, lack the resolution and accuracy of diver surveys in which sponge species and size were mapped on a site. In addition, sources of error for the hyperbolic localization of sound sources (e.g., variations in the speed of sound due to water temperature) would increase the variance around source position estimates.

**Figure 5.**
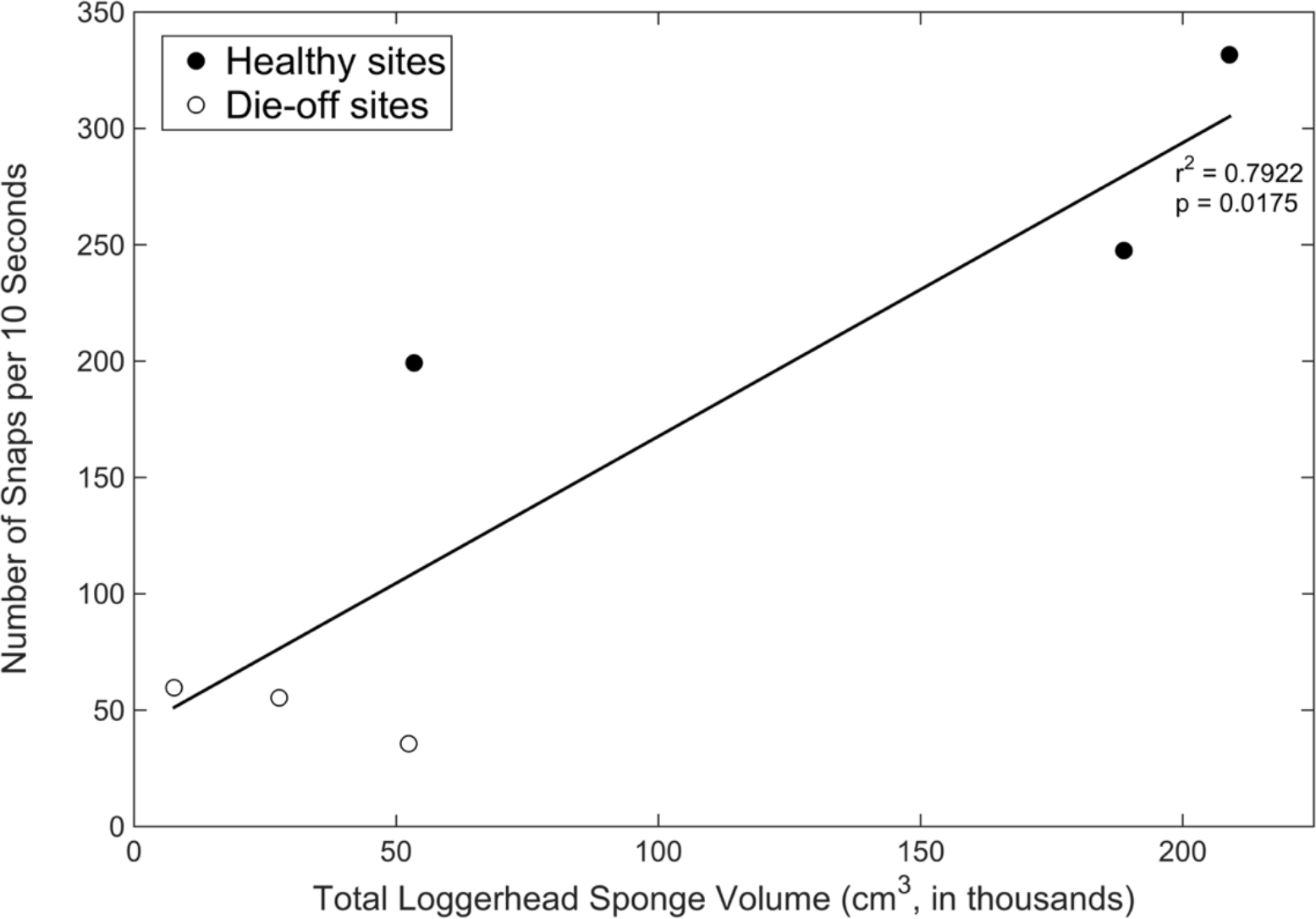
Total loggerhead sponge volume on a given site versus number of snaps per 10 seconds

The use of rapid acoustic monitoring techniques to estimate snapping shrimp abundance and sponge biomass also provides insight into the structure of the habitat in an area. Few studies have linked acoustic metrics to habitat structural complexity and biodiversity (Lammers et al. 2007; Kennedy et al. 2010). Because of the close association between snapping shrimps and sponges, mapping the clusters of snapping shrimp snaps, and hence the sponges in which they dwell, might provide a rapid method to determine the sponge biomass and the structural complexity of that site. Furthering our understanding of how physical and biological characteristics interact with the acoustic environment may provide even more benefits. For example, the larvae of some marine fishes and invertebrates respond to habitat-associated sound cues (Tolimieri et al. 2000, 2004; Simpson et al. 2005, 2008; Stanley et al. 2009, 2011, 2012; Vermeij et al. 2010-among others), and integrating small-scale acoustic and structural variation may help in identifying and predicting patterns of larval recruitment.

In summary, we estimated how the loss of marine sponges from near-shore hard-bottom communities affected snapping shrimp populations by employing remote acoustic recording techniques and distance sampling theory. Areas that suffered sponge die-offs exhibited fewer snapping shrimp snaps and, as a consequence, the estimated snapping shrimp population density and abundance estimates were lower in these areas. This method can hopefully provide rapid assessment of habitat structure and quality, particularly for habitats in which soniferous organisms are associated with structure-forming species.

## 5. Acknowledgements

We are grateful to the many people who provided support and feedback on this project, including C B Butler, L Butler, A J Spadaro, and J Anderson. This research was funded by grants from the NOAA-TNC Community Restoration Program (Award #: GMT-0DU-091512) and Florida SeaGrant (Award #: PD-14-11), and J Butler’s tuition and stipend were provided by the Virginia Modeling and Simulation Center’s Modeling and Simulation Graduate Fellowship. We also thank the Florida Keys National Marine Sanctuary (Permit #: FKNMS-2011-100) for allowing us to conduct research within the sanctuary.

## References Cited

Airoldi, L., Balata, D., Beck, M.W., 2008. The Gray Zone: relationships between habitat loss and marine diversity and their applications in conservation. Journal of Experimental Marine Biology and Ecology 366(1), 8–15.

Au, W.W.L., Banks, K., 1998. The acoustics of the snapping shrimp Synalpheus parneomeris in Kaneohe Bay. Journal of the Acoustical Society of America 103(1), 41–47.

Barlow, J., Taylor, B.L., 2005. Estimates of sperm whale abundance in the northeastern temperate Pacific from a combined acoustic and visual survey. Marine Mammal Science 21(3), 429–445.

Bohnenstiehl, D.R., Lillis, A., Eggleston, D.B., 2016. The Curious Acoustic Behavior of Estuarine Snapping Shrimp: Temporal Patterns of Snapping Shrimp Sound in Sub-Tidal Oyster Reef Habitat. PLoS ONE 11(1).

Bouwma, P.E., Herrnkind, W.F., 2009. Sound production in Caribbean spiny lobster Panulirus argus and its role in escape during predatory attack by Octopus briareus. New Zealand Journal of Marine and Freshwater Research 43(1), 3–13.

Buckland, S.T., Anderson, D.R., Burnham, K.P., Laake, J.L., 2005. Distance sampling. Wiley Online Library.

Buckland, S.T., Anderson, D.R., Burnham, K.P., Laake, J.L., Borchers, D., Thomas, L., 2001. Introduction to distance sampling estimating abundance of biological populations. Oxford University Press, Oxford.

ButlerIV, M.J., Hunt, J.H., Herrnkind, W.F., Childress, M.J., Bertelsen, R., Sharp, W., Matthews, T., Field, J.M., Marshall, H.G., 1995. Cascading disturbances in Florida Bay, USA: cyanobacteria blooms, sponge mortality, and implications for juvenile spiny lobsters Panulirus argus. Marine Ecology Progress Series 129, 119–125.

Butler, J., Stanley, J., Butler IV, M.J., in press. Underwater soundscapes in near-shore tropical habitats and the effects of environmental degradation and habitat restoration. Journal of Experimental Marine Biology and Ecology.

Cato, D., 1976. Ambient sea noise in waters near Australia. The Journal of the Acoustical Society of America 60(2), 320–328.

Cato, D.H., 1980. Some unusual sounds of apparent biological origin responsible for sustained background noise in the Timor Sea. The Journal of the Acoustical Society of America 68(4), 1056–1060.

Cato, D.H., McCauley, R.D., 2002. Australian research in ambient sea noise. Acoustics Australia 30, 1320.

Pew Ocean Commission 2003. America’s Living Oceans: Charting a Course for Sea Change, Arlington, VA.

Cousteau, J.-Y., Malle, L., 1956. The Silent World, pp. 86 minutes.

Diaz, R.J., Rosenberg, R., 2008. Spreading dead zones and consequences for marine ecosystems. Science 321(5891), 926–929.

Dickens, L.C., Goatley, C.H., Tanner, J.K., Bellwood, D.R., 2011. Quantifying relative diver effects in underwater visual censuses. PLoS One6(4), e18965.

Duffy, J.E., 2002. The ecology and evolution of eusociality in sponge-dwelling shrimp. In: Kikuchi, T. (Ed.), Genes, behaviors, and evolution in social insects. University of Hokkaido Press, Sapporo, Japan, pp. 217–252.

Duffy, J.E., Macdonald, K.S., 1999. Colony structure of the social snapping shrimp Synalpheus filidigitus in Belize. Journal of Crustacean Biology 19(2), 283–292.

Halpern, B.S., Walbridge, S., Selkoe, K.A., Kappel, C.V., Micheli, F., D’Agrosa, C., Bruno, J.F., Casey, K.S., Ebert, C., Fox, H.E., 2008. A global map of human impact on marine ecosystems. Science 319(5865), 948–952.

Harris, D., Matias, L., Thomas, L., Harwood, J., Geissler, W.H., 2013. Applying distance sampling to fin whale calls recorded by single seismic instruments in the northeast Atlantic. The Journal of the Acoustical Society of America 134(5), 3522–3535.

Harris, S.A., Shears, N.T., Radford, C.A., 2015. Ecoacoustic indices as proxies for biodiversity on temperate reefs. Methods in Ecology and Evolution.

Jackson, J.B., Kirby, M.X., Berger, W.H., Bjorndal, K.A., Botsford, L.W., Bourque, B.J., Bradbury, R.H., Cooke, R., Erlandson, J., Estes, J.A., 2001. Historical overfishing and the recent collapse of coastal ecosystems. Science 293(5530), 629–637.

Kennedy, E., Holderied, M., Mair, J., Guzman, H., Simpson, S., 2010. Spatial patterns in reef-generated noise relate to habitats and communities: Evidence from a Panamanian case study. Journal of Experimental Marine Biology and Ecology 395(1), 85–92.

Kremen, C., Merenlender, A.M., Murphy, D.D., 1994. Ecological monitoring: a vital need for integrated conservation and development programs in the tropics. Conservation Biology 8(2), 388–397.

Kusel, E.T., Mellinger, D.K., Thomas, L., Marques, T.A., Moretti, D., Ward, J., 2011. Cetacean population density estimation from single fixed sensors using passive acoustics. The Journal of the Acoustical Society of America 129(6), 3610–3622.

Lammers, M.O., Brainard, R.E., Au, W.W., Mooney, T.A., Wong, K.B., 2007. An ecological acoustic recorder (EAR) for long-term monitoring of biological and anthropogenic sounds on coral reefs and other marine habitats. The Journal of the Acoustical Society of America 123(3), 1720–1728.

Lillis, A., Eggleston, D.B., Bohnenstiehl, D.R., 2014. Estuarine soundscapes: distinct acoustic characteristics of oyster reefs compared to soft-bottom habitats. Marine Ecology Progress Series 505, 1–17.

Limburg, K.E., 1999. Estuaries, ecology, and economic decisions: an example of perceptual barriers and challenges to understanding. Ecological Economics 30, 185–188.

Lotze, H.K., Milewski, I., 2004. Two centuries of multiple human impacts and successive changes in a North Atlantic food web. Ecological Applications 14(5), 1428–1447.

Luczkovich, J.J., Mann, D.A., Rountree, R.A., 2008. Passive acoustics as a tool in fisheries science. Transactions of the American Fisheries Society 137(2), 533–541.

Marques, T.A., Thomas, L., Martin, S.W., Mellinger, D.K., Ward, J.A., Moretti, D.J., Harris, D., Tyack, P.L., 2013. Estimating animal population density using passive acoustics. Biological Reviews 88(2), 287–309.

Marques, T.A., Thomas, L., Ward, J., DiMarzio, N., Tyack, P.L., 2009. Estimating cetacean population density using fixed passive acoustic sensors: An example with Blainville’s beaked whales. The Journal of the Acoustical Society of America 125(4), 1982–1994.

McDonald, M., Moore, S., 2002. Calls recorded from North Pacific right whales (Eubalaena japonica) in the eastern Bering Sea. Journal of Cetacean Research and Management 4(3), 261–266.

McDonald, M.A., Fox, C.G., 1999. Passive acoustic methods applied to fin whale population density estimation. The Journal of the Acoustical Society of America 105(5), 2643–2651.

McWilliam, J.N., Hawkins, A.D., 2013. A comparison of inshore marine soundscapes. Journal of Experimental Marine Biology and Ecology 446, 166–176.

Mellinger, D., Barlow, J., 2003. Future directions for acoustic marine mammal surveys: stock assessment and habitat use. Pacific Marine Environment Laboratory, Report of a workshop held in La Jolla, CA November 20-22, 2002. Technical contribution No. 2557.

Mellinger, D.K., Stafford, K.M., Fox, C.G., 2004. Seasonal occurrence of sperm whale (Physeter macrocephalus) sounds in the Gulf of Alaska, 1999-2001. Marine Mammal Science 20(1), 48–62.

Mellinger, D.K., Stafford, K.M., Moore, S., Dziak, R.P., Matsumoto, H., 2007. Fixed passive acoustic observation methods for cetaceans. Oceanography 20(4), 36.

Moore, S.E., Stafford, K.M., Mellinger, D.K., Hildebrand, J.A., 2006. Listening for large whales in the offshore waters of Alaska. BioScience 56(1), 49–55.

Moretti, D., DiMarzio, N., Morrissey, R., Ward, J., Jarvis, S., 2006. Estimating the density of Blainville’s beaked whale (Mesoplodon densirostris) in the Tongue of the Ocean (TOTO) using passive acoustics. MTS/IEEE OCEANS.

Myrberg Jr, A.A., 1981. Sound communication and interception in fishes. In: Tavolga, W.N., Popper, A.N., Fay, R.R. (Eds.), Hearing and sound communication in fishes. Springer, New York, pp. 395–426.

Pijanowski, B.C., Villanueva-Rivera, K.J., Dumyahn, S.L., Farina, A., Krause, B.L., Napoletano, B.M., Gage, S.H., Pieretti, N., 2011. Soundscape Ecology: The Science of Sound in the Landscape. BioScience 61(3), 203–216.

Pile, A., Patterson, M., Witman, J., 1996. In situ grazing on plankton < 10 μm by the boreal sponge Mycale lingua. Marine Ecology Progress Series 141, 95–102.

Radford, C., Jeffs, A., Tindle, C., Montgomery, J.C., 2008. Resonating sea urchin skeletons create coastal choruses. Marine Ecology Progress Series 362, 37–43.

Radford, C., Stanley, J., Tindle, C., Montgomery, J.C., Jeffs, A.G., 2010. Localised coastal habitats have distinct underwater sound signatures. Marine Ecology Progress Series 401, 21–29.

Radford, C.A., Jeffs, A.G., Tindle, C.T., Montgomery, J.C., 2008. Temporal patterns in ambient noise of biological origin from a shallow water temperate reef. Oecologia 156(4), 921–929.

Reiswig, H., 1990. In situ feeding in two shallow-water Hexactinellid sponges. New Perspectives in Sponge Biology. Smithsonian Institution Press, Washington, DC, 504–510.

Scharer, M.T., Nemeth, M.I., Rowell, T.J., Appeldoorn, R.S., 2014. Sounds associated with the reproductive behavior of the black grouper (Mycteroperca bonaci). Marine Biology 161(1), 141–147.

Simpson, S.D., Meekan, M., Montgomery, J., McCauley, R., Jeffs, A., 2005. Homeward sound. Science 308(5719), 221–221.

Simpson, S.D., Meekan, M.G., Jeffs, A., Montgomery, J., McCauley, R., 2008. Settlement-stage coral reef fish prefer the higher-frequency invertebrate-generated audible component of reef noise. Animal Behavior 75, 1861–1868.

Sirovic, A., Hildebrand, J.A., Wiggins, S.M., McDonald, M.A., Moore, S.E., Thiele, D., 2004. Seasonality of blue and fin whale calls and the influence of sea ice in the Western Antarctic Peninsula. Deep Sea Research Part II: Topical Studies in Oceanography 51(17), 2327–2344.

Sirovic, A., Hildebrand, J.A., Wiggins, S.M., Thiele, D., 2009. Blue and fin whale acoustic presence around Antarctica during 2003 and 2004. Marine Mammal Science 25(1), 125–136.

Solan, M., Cardinale, B.J., Downing, A.L., Engelhardt, K.A., Ruesink, J.L., Srivastava, D.S., 2004. Extinction and ecosystem function in the marine benthos. Science 306(5699), 1177–1180.

Spiesberger, J.L., 1999. Locating animals from their sounds and tomography of the atmosphere: Experimental demonstration. The Journal of the Acoustical Society of America 106(2), 837–846.

Spiesberger, J.L., 2001. Hyperbolic location errors due to insufficient numbers of receivers. The Journal of the Acoustical Society of America 109(6), 3076–3079.

Spiesberger, J.L., Fristrup, K.M., 1990. Passive localization of calling animals and sensing of their acoustic environment using acoustic tomography. American Naturalist, 107–153.

Staaterman, E., Paris, C.B., Kough, A.S., 2014. First evidence of fish larvae producing sounds. Biology Letters 10(10), 20140643.

Stanley, J.A., Radford, C.A., Jeffs, A.G., 2010. Induction of settlement in crab megalopae by ambient underwater reef sound. Behavioral Ecology 21(1), 113–120.

Stanley, J.A., Radford, C.A., Jeffs, A.G., 2011. Behavioural response thresholds in New Zealand crab megalopae to ambient underwater sound. PLoS One 6(12), e28572–e28572.

Stanley, J.A., Radford, C.A., Jeffs, A.G., 2012. Location, location, location: finding a suitable home among the noise. Proceedings of the Royal Society B 279, 3622–3631.

Stevely, J.M., Sweat, D.E., Bert, T.M., Sim-Smith, C., Kelly, M., 2011. Sponge mortality at Marathon and Long Key, Florida: patterns of species response and population recovery. Proceedings of the 63rd Gulf and Caribbean Fisheries Institute, 384–400.

Suchanek, T.H., 1994. Temperate coastal marine communities: biodiversity and threats. American Zoologist 34(1), 100–114.

Tait, R.I., 1962. The evening chorus: a biological noise investigation. Naval Research Laboratory, MHNZ Dockyad, Aukland.

Thomas, L., Buckland, S.T., Rexstad, E.A., Laake, J.L., Strindberg, S., Hedley, S.L., Bishop, J.R., Marques, T.A., Burnham, K.P., 2010. Distance software: design and analysis of distance sampling surveys for estimating population size. Journal of Applied Ecology 47(1), 5–14.

Willis, T.J., 2001. Visual census methods underestimate density and diversity of cryptic reef fishes. Journal of Fish Biology 59, 1408–1411.

Tolimieri, N., Haine, O., Jeffs, A., McCauley, R., Montgomery, J., 2004. Directional orientation of Pomacentrid larvae to ambient reef sound. Coral Reefs 23, 184–191.

Tolimieri, N., Jeffs, A., Montgomery, J.C., 2000. Ambient sound as a cue for navigation by the pelagic larvae of reef fishes. Marine Ecology Progress Series 207, 219–224.

Urick, R.J., 1983. Principles of underwater sound. McGraw-Hill Book Company, New York.

Valiela, I., Bowen, J.L., York, J.K., 2001. Mangrove Forests: One of the World’s Threatened Major Tropical Environments At least 35% of the area of mangrove forests has been lost in the past two decades, losses that exceed those for tropical rain forests and coral reefs, two other well-known threatened environments. Bioscience 51(10), 807–815.

Van Opzeeland, I., Kindermann, L., Boebel, O., Van Parijs, S., 2008. Insights into the acoustic behaviour of polar Pinnipeds-current knowledge and emerging techniques of study. In: Weber, E., Krause, L. (Eds.), Animal Behaviour: New Research. Nova Science Publishers, Hauppage, NY.

Vermeij, M.J., Marhaver, K.L., Huijbers, C.M., Nagelkerken, I., Simpson, S.D., 2010. Coral larvae move toward reef sounds. PLOS One 5(5).

Versluis, M., Schmitz, B., von der Heydt, A., Lohse, D., 2000. How snapping shrimp snap: through cavitating bubbles. Science 289(5487), 2114–2117.

Vitousek, P.M., Mooney, H.A., Lunchenco, J., Melillo, J.M., 1997. Human domination of the Earth’s ecosystems. Science 277, 494–499.

Watanabe, M. Sekine, E. Hamada, M. Ukita and T. Imai. 2002. Monitoring of shallow sea environment by using snapping shrimps. Water Science & Technology 46(1): 419–424.

Waycott, M., Duarte, C.M., Carruthers, T.J., Orth, R.J., Dennison, W.C., Olyarnik, S., Calladine, A., Fourqurean, J.W., Heck, K.L., Hughes, A.R., 2009. Accelerating loss of seagrasses across the globe threatens coastal ecosystems. Proceedings of the National Academy of Sciences 106(30), 12377–12381.

Worm, B., Barbier, E.B., Beaumont, N., Duffy, J.E., Folke, C., Halpern, B.S., Jackson, J.B., Lotze, H.K., Micheli, F., Palumbi, S.R., 2006. Impacts of biodiversity loss on ocean ecosystem services. Science 314(5800), 787–790.

